# *Cis*-nonProline Peptides: Genuine Occurrences and their Functional Roles

**DOI:** 10.1101/324517

**Authors:** Jane S. Richardson, Lizbeth L. Videau, Christopher J. Williams, Bradley J. Hintze, Steven M. Lewis, David C. Richardson

## Abstract

While *cis* peptides preceding proline can occur about 5% of the time, *cis* peptides preceding any other residue (“*cis*-nonPro” peptides) are an extremely rare feature in protein structures, of considerable importance for two opposite reasons. On one hand, their genuine occurrences are mostly found at sites critical to biological function, from the active sites of carbohydrate enzymes to rare adjacent-residue disulfide bonds. On the other hand, a *cis*-nonPro can easily be misfit into weak or ambiguous electron density, which led to a high incidence of unjustified *cis*-nonPro over the 2006-2015 decade. This paper uses high-resolution crystallographic data and especially stringent quality-filtering at the residue level to identify genuine occurrences of *cis*-nonPro, and to survey both individual examples and broad patterns of their functionality.

We explain the procedure developed to identify genuine *cis*-nonPro examples with almost no false positives. We then survey a large sample of the varied functional roles and structural contexts of *cis*-nonPro, including the uses of specific amino acids for particular purposes. We emphasize aspects not previously covered: that *cis*-nonPro always (except for vicinal disulfides) occur in highly ordered structure, and especially the great concentration of occurrence in proteins that process or bind carbohydrates (identified by occurrence on the CAZy website).

## Introduction

Peptide bonds in protein structures have a partial double-bond character, which keeps them fairly close to planar. The majority adopt the *trans* backbone conformation, with the ω dihedral angle near 180°, while some are *cis* with ω near 0°. Peptides preceding prolines have somewhat less unfavorable energy difference and a lower barrier to *cis/trans* rotation (reviewed in Pal et al., 1999), and the reported occurrence frequency of *cis*-Pro has converged over the years to a bit over 5% (Stewart et al., 1990; MacArthur et al., 1991; Jabs et al., 1999), more common in β-sheet proteins than in helical ones. *Cis*-Pro play structural roles, as exposed turns or to accommodate tightly packed regions in the interior. They also play static functional roles such as positioning active-site residues in carbonic anhydrase and in pectate lyase (Videau et al., 2004). Such roles give them rather high conservation within a protein family (Lorenzen et al., 2005). On the dynamic side, a need to achieve this rarer isomer produces a slow step in the protein folding process (Brandts et al., 1975; Schmid et al., 1992), and there are well-studied proline isomerase enzymes which aid in that step (Wawra et al., 2006). The slow *cis/trans* conversion has been exploited by evolution for its suitability in biological timing circuits (Lu et al., 2007).

In contrast, genuine *cis* peptides preceding non-proline residues are extremely scarce. Historically, the first examples were three *cis*-nonPro near the active site of carboxypeptidase A (Rees et al., 1981) and the Gly-*cis*-Gly peptide that helps bind both NADPH and inhibitors in dihydrofolate reductase (Bolin et al., 1982). Figure 1 shows that site at 1.09A resolution in the later 1kms human DHFR (Klon et al., 2002).

**Figure 1.**
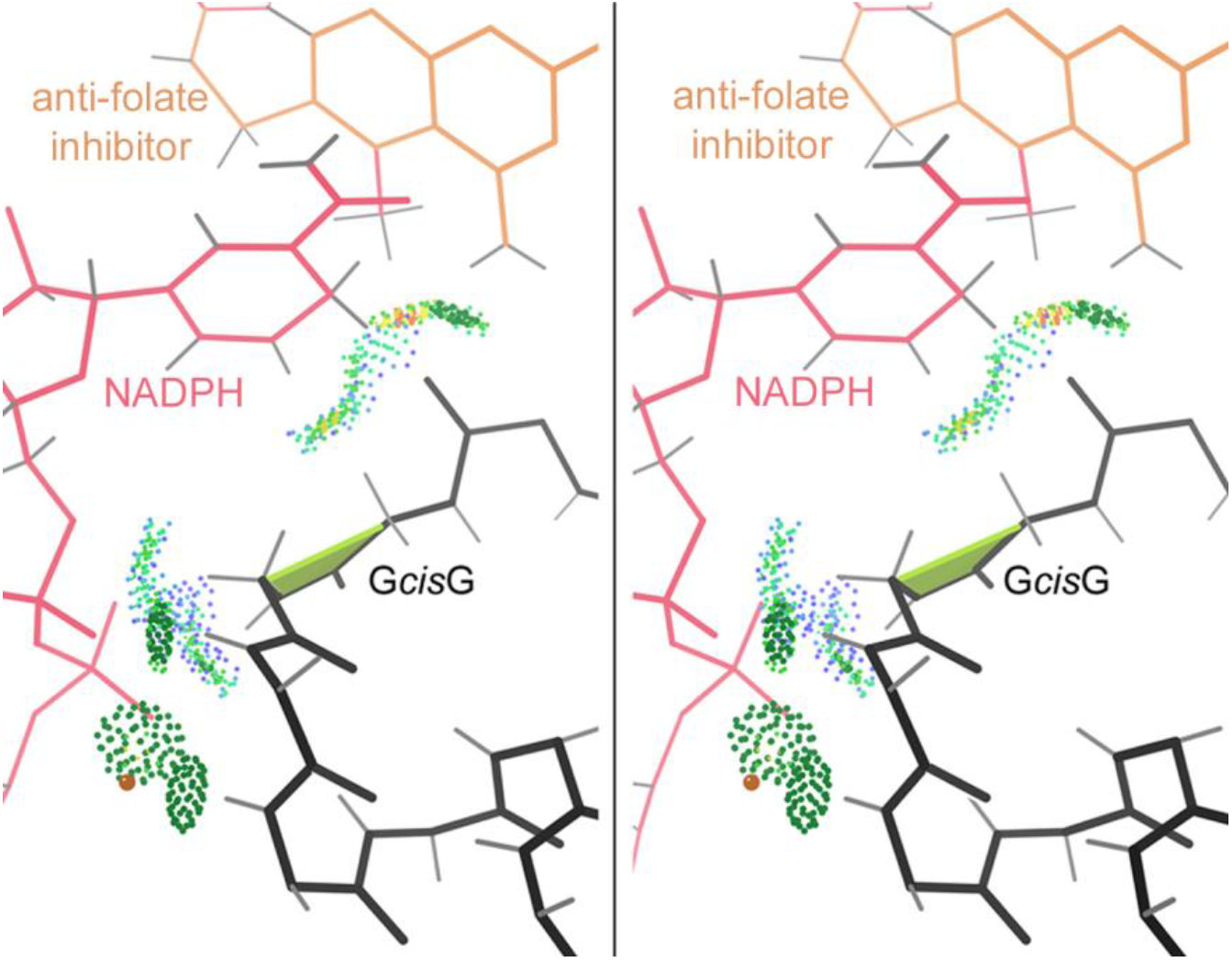
Walleye stereo of one of the first *cis*-nonPro identified, in dihydrofolate reductase, as shown in the later 1.09A resolution 1kms. The *cis* peptide is marked by a lime trapezoid, while the all-atom contacts are shown as pairs of dots for van der Waals contacts and darker green pillows for H-bonds.

Specific subcategories of genuine *cis*-nonPro (Richardson et al., 2017) and of systematically incorrect *cis*-nonPro (Richardson et al., 2022; Williams et al., 2022a, 2022b) have been described.

## Results

### Reliability filtering for genuine occurrences

Any search for an unusual conformation needs quality filtering, at the residue level as well as the file level, in order to achieve a robust structural-bioinformatics treatment (Williams et al., 2022a). Even more rigorous filtering is required for *cis*-nonPro peptides, given the epidemic of over-use in experimental structures, in poor density at all resolutions (Croll 2015; Williams et al., 2015). Therefore, as well as steric and geometrical criteria, for this analysis we also use criteria based on electron density quality (see Methods section), modified from those first used in our latest sidechain rotamer analysis (Hintze et al., 2016). This added filter will reject some genuine cases, but has been shown to accept extremely few incorrect cases, and so is effective for choosing a list of reliably correct *cis*-nonPro examples for evaluating factors that affect occurrence preferences and for studying the real functional roles of this conformation.

The resulting reliability-filtered list contains 437 *cis*-nonPro peptides in 378 different protein chains, compiled from the Top8000 reference dataset of 6765 protein chains at < 2.0A resolution, < 50% sequence homology, with deposited diffraction data, and additional residue-level filters. This *cis*-nonPro list, with annotations, is given in Table S1 of the supplementary material. It forms the primary basis of the analyses in this paper, with a few exceptions that investigate relationships or conservation across a wider range.

### Preferential occurrence of *cis*-nonPro in carbohydrate-active proteins

It has been known for quite some time that a number of carbohydrate enzymes make use of *cis*-nonPro conformations (Jabs et al., 1999), and we are now in a position to judge whether this apparent connection is significant. Our source for automated identification of carbohydrate-processing or binding proteins is inclusion on the CAZy web site (Carbohydrate Active enZymes; http://www.cazy.org; Lombard et al., 2014). 99 of our 378 distinct protein chains with a genuine *cis*-nonPro (26%) are in CAZy, while only 6% of the 6765 total chains in our reference dataset (the Top8000_50%_SF) are in CAZy. Carbohydrate-active protein chains are thus over-represented by a factor of >4 in this list. Of the 439 reliably genuine *cis*-nonPro peptides, 144 examples (33%) occur in proteins that are on CAZy. Another way of expressing this contrast is that 76% (33/43) of the protein chains with more than one authenticated *cis*-nonPro peptide are in CAZy. Those 33 chains are only 0.49% of the reference dataset, but they contain 77 (17.5%) of the *cis*-nonPro peptides, an over-representation of 36-fold. Thus carbohydrate-active enzymes are very much more likely than other proteins to have more than a single genuine *cis*-nonPro per chain, as well as much more likely to have any at all.

Of the 43 protein chains with > 1 authenticated *cis*-nonPro, 15 (34%) are chitinases or chitinase-like: 9 of them have three *cis*-nonPros and 6 have two (see Table S1). We found a later-deposited protein chain with 4 genuine *cis*-nonPro, also a chitinase: 3wd0. Chitinases are remarkably widespread and sequence-diverse throughout phyla, occurring in bacteria, fungi, insects, plants, animals, and archaea (Adrangi and Faramarzi 2013; Nakamura et al., 2007), providing either nutritional use, protection against, or other uses of chitin-containing arthropod and especially fungal organisms. Their diversity explains how so many of them can occur in a dataset with < 50% sequence homology.

The positions of *cis*-nonPro peptides in a sample of chitinases are shown in a cross-sectional slice in Figure 2. All family 18 chitinases have a Trp-*cis*-X *cis*-nonPro at the C-terminal end of β strand 8 in their TIM-barrel fold, where the Trp is well packed and seems to act by positioning the backbone and residue X, which is usually the catalytic acid. In the triple *cis*-nonPro chitinases, the other two well-conserved *cis*-nonPros are at or near the ends of β strands 2 and 4, and are involved in substrate binding (Terwissa van Scheltinga 1996).

**Figure 2.**
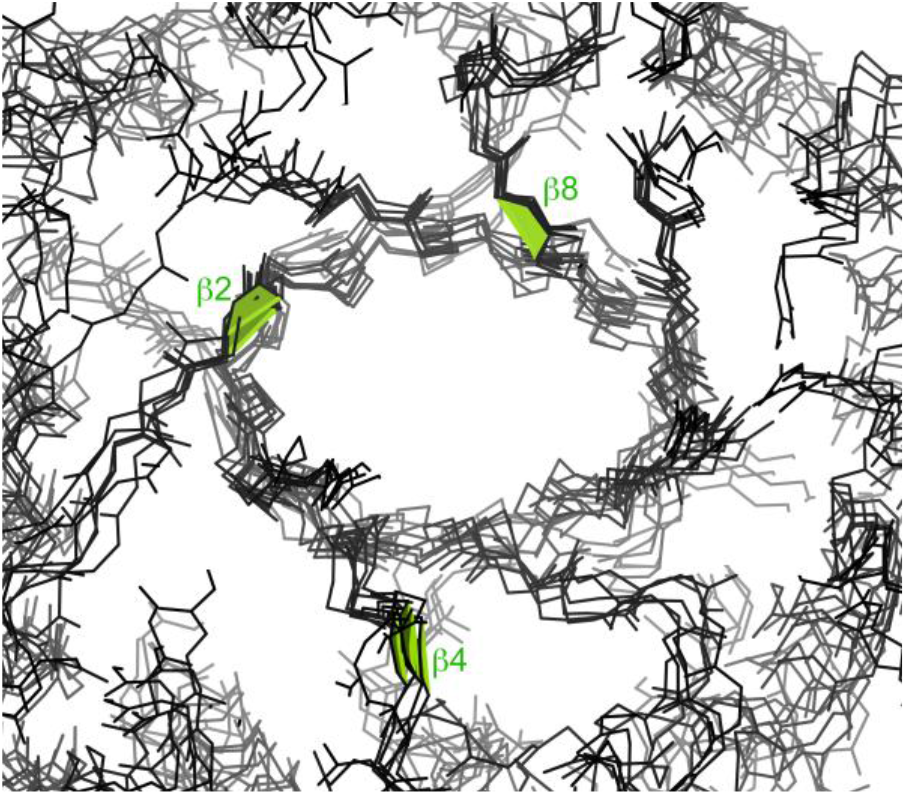
Six chitinase TIM barrels superimposed, showing the completely conserved *cis*-nonPros at the end of β strand 8 (all Trp-*cis*-X) and β strand 2, and the less conserved ones at the end of β strand 4. They are 3n11 *Bacillus cereus*, missing β4 (Hsieh et al., 2010), 1goi *Serratia marcescens* (Kolstad et al., 2002), 1w9p *Aspergillus fumigatus* (Hurtado-Guerrero et al., 2007), 2uy2 *Saccharomyces cerevisiae*, missing β4 (Rao et al., 2005), 3fy1 acidic mammalian (Olland et al., 2009), and 3alf *Nicotiana tabacum* (Ohnuma et al., 2011), at 1.2 – 1.7A resolution.

Besides the chitinases, chains with three *cis*-nonPro peptides include two β-galactosidases (1yq2 and 3cmg), in CAZy, and three carboxypeptidases (2piy, 3d4u, and 3i1u), not in CAZy. The chains with two *cis*-nonPro peptides include a wider variety of functions, 14 in and 6 out of CAZy.

The *cis*-nonPro-containing carbohydrate enzymes are unevenly distributed across the many structure/function families defined in CAZy. There are 17 *cis*-nonPro protein chains in family GH18 (glycoside hydrolase 18): the 15 chitinases and 2 xylanase inhibitors. There are 13 in family GH5, mostly mannanases, 11 in family GH1, mostly β-glucosidases, 8 in family GH10, all xylanases, 5 in family GH2, and 5 in different PL families (polysaccharide lyases). In contrast, the remaining 41 CAZy *cis*-nonPro chains are spread across 34 different families (see Table S1). The xylanases present an interesting combination of conservation versus divergence. The 8 in family GH18 all use a His-*cis*-Thr *cis* peptide at the end of β-strand 3 of their TIM barrels. The 3 xylanases in other GH families each have a different *cis*-peptide sequence, located at the end of β-strand 1 (2ddx), β-strand 4 (2y8k), or β-strand 5 (1nof) of their TIM barrels.

Lectins, or CBMs (Carbohydrate-Binding Modules), are listed currently on CAZy only when they are part of a protein that also includes a carbohydrate enzyme, either as a separate domain or a separate chain. Therefore, we did not have an automated way of identifying all of them uniformly. However, anecdotally, lectins such as the prototypical concanavalin A are fairly well represented on the *cis*-nonPro list (14 of them), but not as over-represented as the carbohydrate enzymes. The CBMs mostly have all-β antiparallel folds. If the 14 lectins in our list, plus the 47 whose names indicate action on a carbohydrate, are added to the 99 enzymes that are in CAZy, then 160 of the 378 *cis*-nonPro-containing chains, or 42%, are carbohydrate related. Since we made these additional assignments mainly from the titles and abstracts on the PDB site, they are not necessarily complete or correct, but even as an approximation this means that the predominance of carbohydrate relatedness is even stronger than the pure CAZy survey indicates.

The non-CAZy *cis*-nonPro proteins include 17 phosphoribosyl transferases, another highly sequence-diverse group (the PRT family) with only a 13-residue conserved sequence signature recognized. They share an essential *cis*-nonPro conserved in conformation but not in sequence, on the “PPi loop”. As shown in Figure 3 for the 1.05A 1fsg PRT structure (Heroux et al., 2000) the backbone of the *cis*-nonPro and its adjacent peptides are used to bind a phosphate oxygen of PRPP and, through waters, a Mg ion (Sinha et al., 2001). In at least some PRTs, a *trans* to *cis* change accompanies substrate binding (Shi et al., 2002).

**Figure 3.**
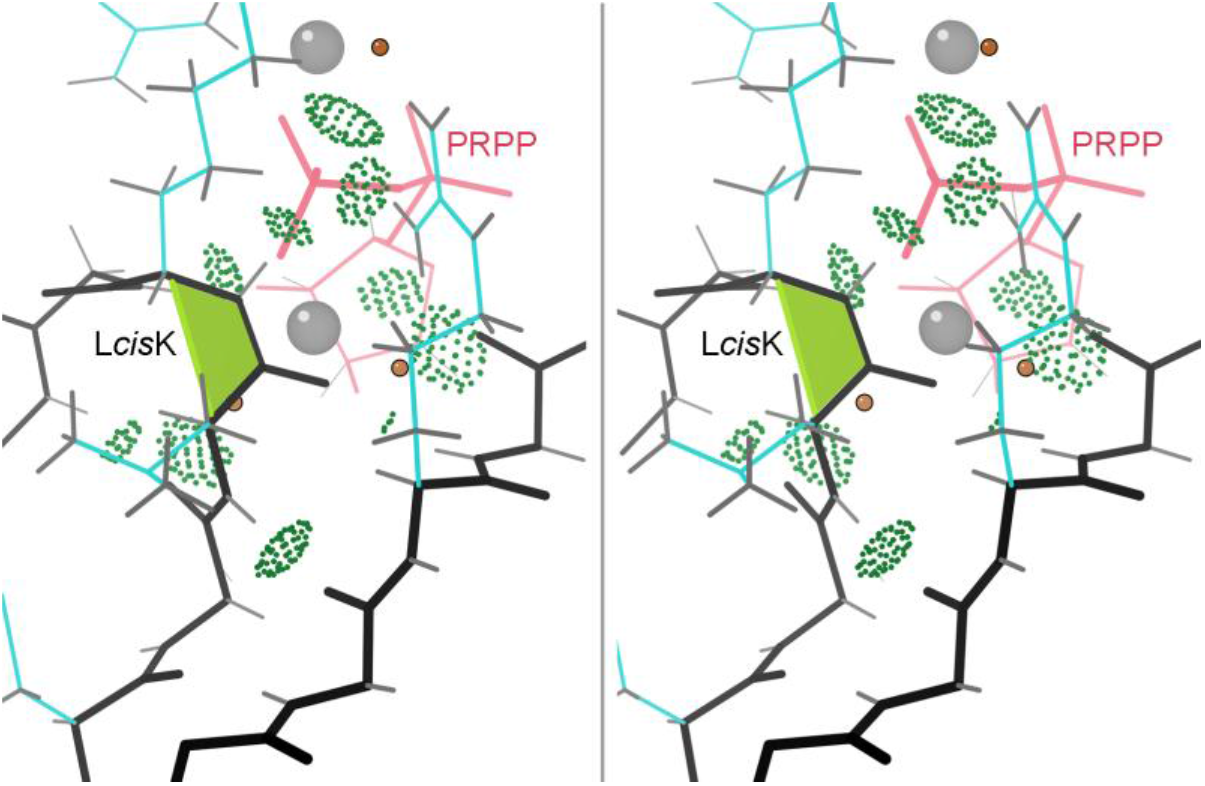
Walleye stereo of 1fsg, a 1.05A phosphoribosyl transferase (PRT) enzyme, showing the many H-bonds between the *cis*-nonPro and the substrate (in pink). Grey balls are Mg ions.

### Preferential occurrence of *cis*-nonPro peptides in TIM-barrel proteins

As well as their enzymatic activity on carbohydrates, another characteristic shared by many of the *cis*-nonPro containing protein chains is a (β/α)_8_ TIM barrel fold. Over half the relevant domains of the 99 CAZy proteins are TIM barrels, plus five (β/α)_8_ domains in phosphodiesterases. However, in the non-CAZy chains, TIM barrels are not especially over-represented, and indeed most TIM barrels (including the eponymous triose phosphate isomerase itself) do not include any *cis*-nonPro peptides. It seems, then, that the strong preference for (β/α)_8_ folds is probably not an independent factor, but is in some way correlated with presence in the carbohydrate-processing system.

The more generic preference for β structure, and for location at or near the C-terminal end of a β strand, does hold for non-CAZy as well as CAZy proteins.

### 5-dimensional ϕ,ψ,ω,ϕ,ψ local conformational and sequence preferences

A useful study tool is to plot datapoints for the genuine *cis*-nonPro examples in the 5-dimensional space of ϕ1, ψ1, ω, ϕ2, ψ2 selectable by their first and second amino-acid identities.

These plots can be viewed 3 dimensions at a time in the Mage interactive graphics program (Richardson et al., 2001). Not surprisingly, there is little spread from 0° in ω, but somewhat more than in trans conformation (see Supplementary table). ψ1, ϕ2 provides the most diagnostic projection, shown in Figure 4a for all *cis*-nonPro and in Figure 4b for selected clusters.

**Figure 4.**
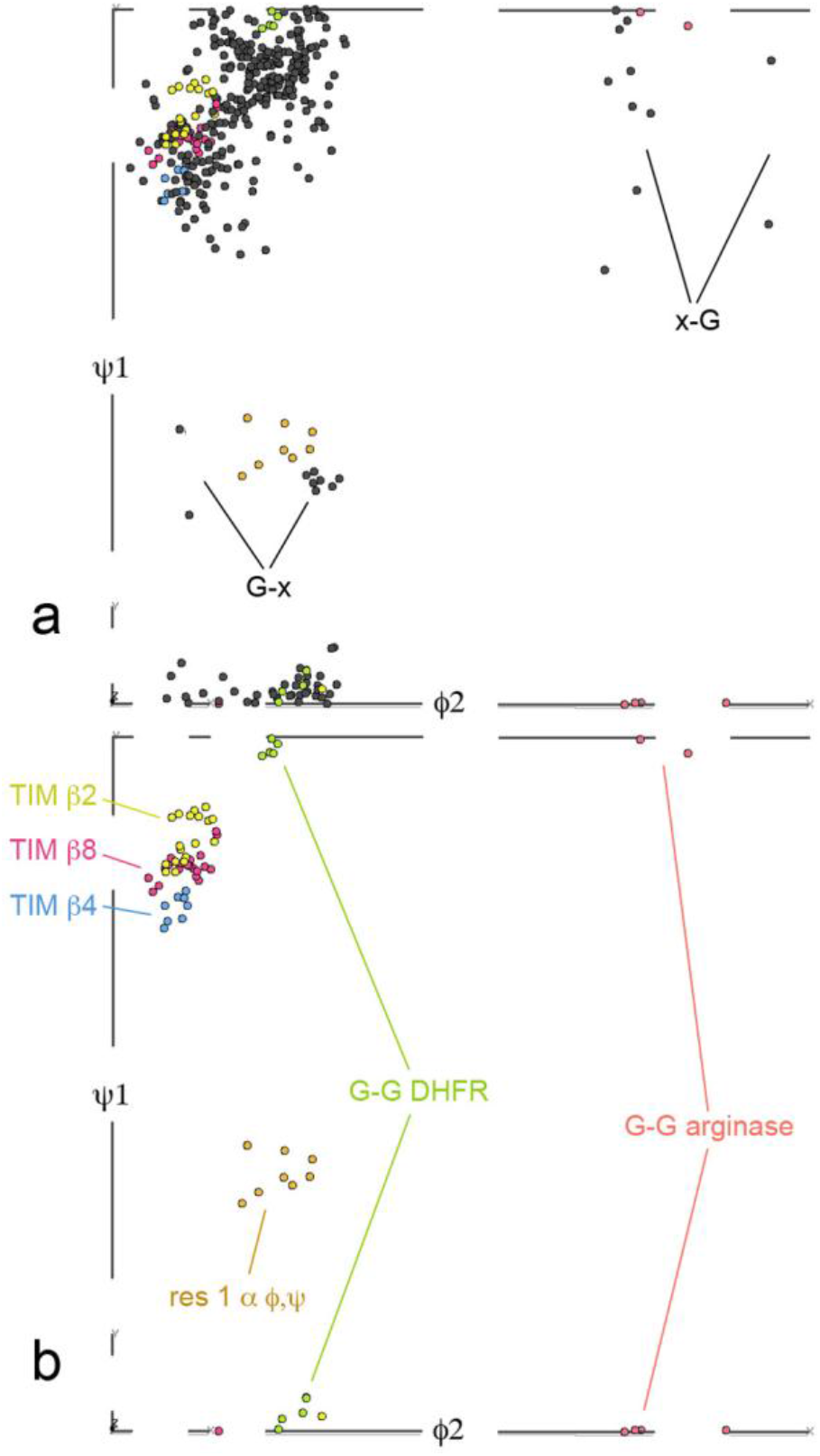
ψ1, ϕ2 plots, taken as the most diagnostic 2-dimensional projections from 5-dimensional plots. Part 4a includes all *cis*-nonPro, and shows the very strong preference for extended structure and the X-*cis*-G, G-*cis*-X and first-residue alpha that are the only exceptions. Part 4b shows the 3 TIM barrel clusters, the G-*cis*-G DHFR and arginase clusters, and the first-residue alpha clusters discussed in the text.

### Trp-X *cis*-nonPro

Trp and Cys are the least frequent amino acids in globular proteins (∼1.3%). However, there are 49 Trp-*cis*-X *cis*-nonPro in our dataset of 437, or 11%, an enrichment by a factor of 8, the highest ratio for any amino acid type. The preference is asymmetrical, with only 13 X-*cis*-Trp and no Trp-*cis*-Trp. This seems to be a functional rather than an energetic preference, since nearly all are on β strand 8 of a TIM-barrel carbohydrate-cleaving CAZy enzyme (14 chitinases, 7 glucosidases, 6 mannanases, 5 endoglucanase cellulases, and 6 other types in our dataset). Those ϕ,ψ values are all closely clustered in all 5 dimensions near the TIM β8 points of Figure 4b, and the Trp sidechain adopts a **t-90** rotamer that lets it usually make some contact with the X sidechain (usually the active-site residue) and often with the adjacent β strand (see Figure 3b). Trp-*cis*-Ser is the most common sequence, followed by Trp-*cis*-Glu.

Five more Trp-*cis*-X *cis*-nonPros are Trp-*cis*-Thr in non-CAZy glycerol-phosphodiester phosphodiesterases, at the end of β−strand 8 of 8-stranded TIM-like barrels. Only one Trp-*cis*-X is in a protein with no carbohydrate relationship at all: 1qgu nitrogenase.

### First-residue helical ϕ,ψ

A *cis* peptide cannot occur inside a regular helix, since it would break the helix. Even single-residue helical ϕ,ψ values are very unusual for *cis*-nonPro, especially in the first amino-acid position. This loose cluster of 8 examples, however, turned out to have nothing else in common. Most examples are some sort of turn, loop, or corner, while 3pb6 Asp-*cis*-Ser makes a bulge in a helix and ligands the Zn. Two are especially unusual and interesting. 2eab A His759-*cis*-Ala-*cis*-Pro761 (Nagae et al., 2007) is an unprecedented type of tight turn formed by successive *cis*-nonPro and *cis*-Pro peptides, confirmed by definitive 1.12A electron density. It has no evident functionality, and we suspect it might not be conserved, but could not tell at the time of the Top8000’s construction because its crystal structure defined a new family (GH95) of fucosidases. Since then, there have been 4 further structures done in the GH95 family for other species (7kmq, 2rdy, 4ufc, and 7znz), all of which have the structure His-*cis*-Pro-*cis*-Pro, with the much commoner *cis*-Pro in both positions.

The one α-first case that has definite biological function is 3hhs chain A Glu-*cis*-Ala354 (Figure 5) and chain B Glu-*cis*-Ser352 (Li et al., 2009). They are conserved in related prophenyl-oxidase enzymes. The *cis*-nonPro forms the C-cap of one helix and makes two backbone H-bonds to the first turn of the next helix, separated by a short, meandering loop. The protruding *cis* peptide puts a bend of about 30° between the two helices. Glu 352 of the *cis*-nonPro stabilizes one of the 6 His ligands to the bi-nuclear Cu site, and the second helix contributes two more His ligands.

**Figure 5.**
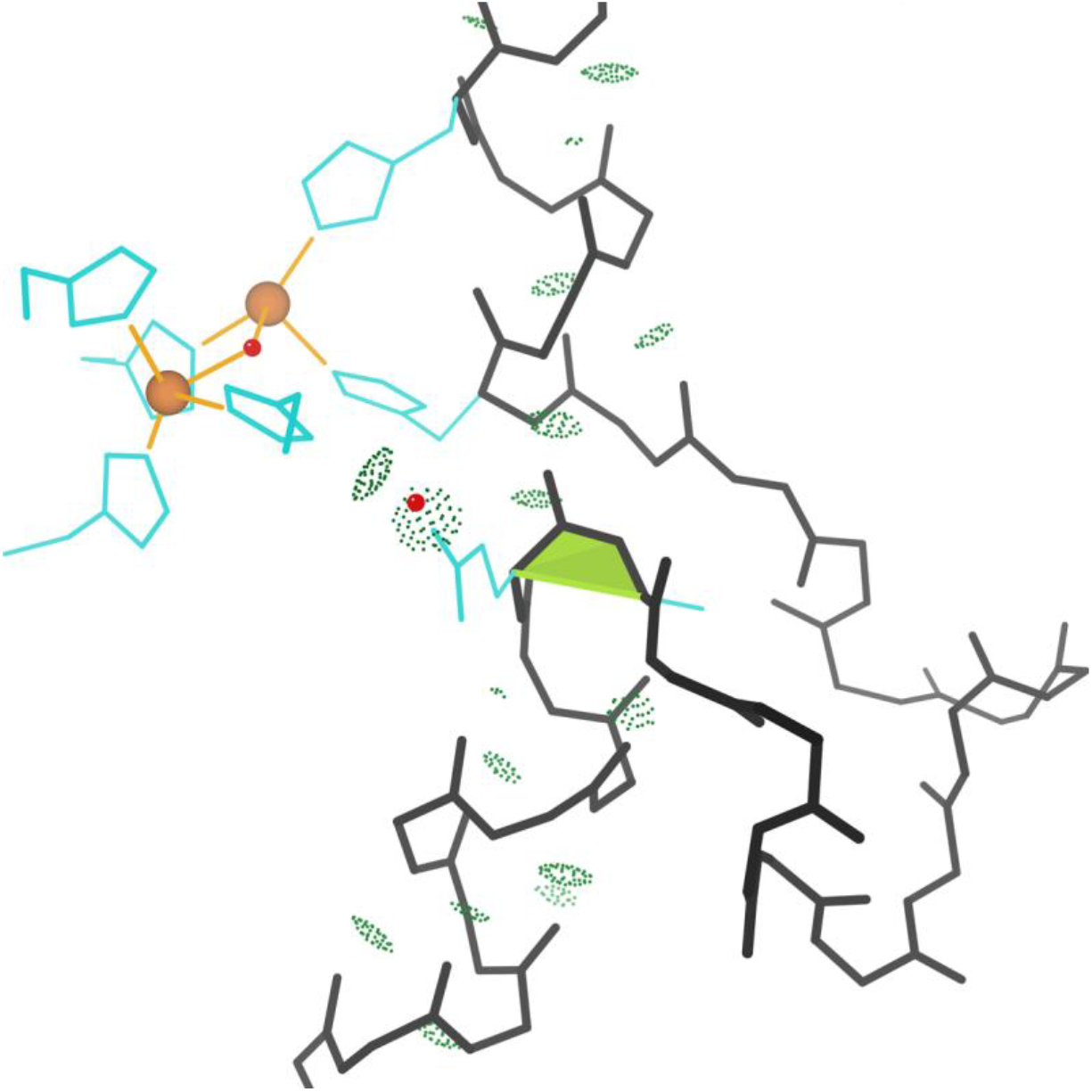
The biologically relevant first-residue-alpha *cis*-nonPro in 3hhs 1.97A prophenyl-oxidase. The Glu is positioned to make an H-bond through water (small red sphere) to a ligand of the bi-nuclear Cu site (large copper-colored spheres), while 2 other ligands are provided by the second helix. H-bonds are shown as green pillows of dots.

### DHFRs & other Gly-*cis*-Gly

Gly is enriched generally by about 3-fold in *cis*-nonPro, at least partly because a Gly-containing peptide is easier to fit incorrectly as *cis* and thus their numbers are inflated in unfiltered data. There is also a functional preference for Gly in some contexts, especially for Gly-*cis*-Gly. Expectation for Gly-*cis*-Gly would be 3.4 examples in our dataset, and there are actually 19. 10 of them are in dihydrofolate reductases (DHFRs) as a conserved and essential feature of the active site. Figure 1 illustrates the local conformation and interactions in the classic 1.09A 1kms *Lactobacillus casei* DHFR Gly-*cis*-Gly, showing how it is a major factor in binding both the NADPH cofactor and the methotrexate inhibitor, using H-bonds from the *cis* and surrounding peptides and van der Waals contacts from the essential Gly H “sidechains”.

These features are all conserved in DHFRs, as shown even more cleanly in the later structures of our dataset such as the 1.09A 1kms human DHFR.

The other Gly-*cis*-Gly *cis*-nonPros are quite varied, including 3 in arginase-superfamily enzymes and 3 in carbohydrate enzymes with β-helix folds. Their conformational clusters are in orange in Figure 4b, quite distinct from the DHFR cluster.

There are 59 Gly-*cis*-X and only 15 X-*cis*-Gly. We do not know of a specific reason for this, but certainly the relationship of the glycines to the rest of the structure is quite different in the two positions.

### Asp-*cis*-Asp

Most of the nine Asp-*cis*-Asp *cis*-nonPro peptides arrange their two sidechain carboxyl groups to form one side of a divalent-cation binding site, where the other side is bound by substrate. Figure 6 shows an example from 2xjp at 0.85A resolution, with bound Ca^++^ and mannose.

**Figure 6.**
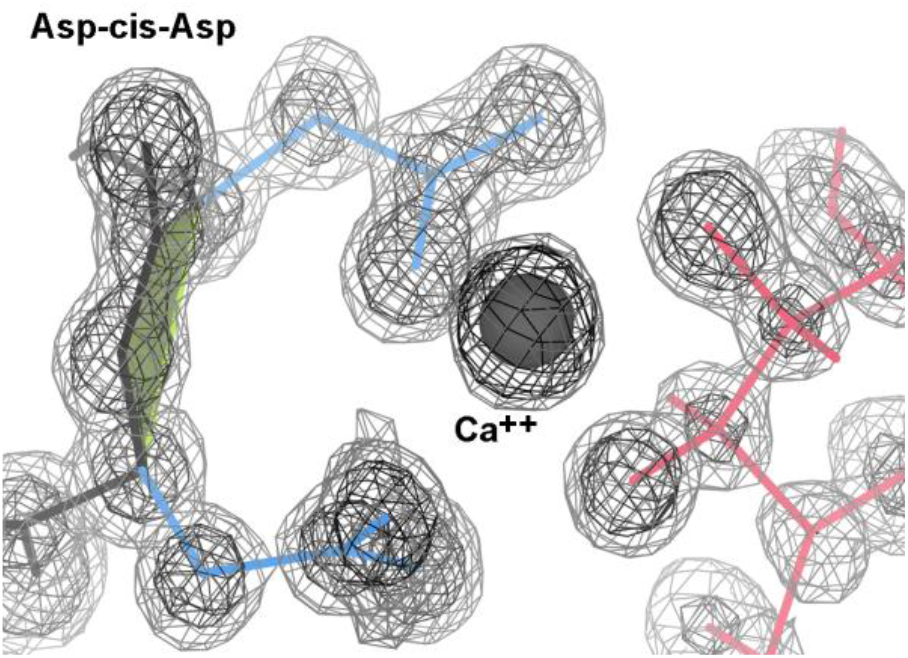
The 2xjp structure of flocculin at 0.85A, where both Asps bind the Ca^++^, which binds the mannose (Veelders et al., 2010). Electron density contours are at 1.2 and 3.0σ.

This flocculin enzyme in *Candida albicans* promotes the self-aggregation called flocculation, one of the steps in brewing beer.

Four other Asp-*cis*-Asp examples bind the Zn^++^ (3ife, 3iib, 3pfe) or Mn^++^ (2pok) at the active site of a metallopeptidase. In the 2jdi F1-ATPase α-chain, just Asp269 binds the Mg^++^ through a water.

2rb7 is a metallopeptidase from *Desulfovibrio desulfuricans* G2 with no metal binding in the crystal and no publication (JCSG 2007), but the Asp-*cis*-Asp is part of a suggestive cluster of 3 Asp, 2 Glu, and 2 His deep in a cleft. At 30-40% homology, there is a new pdb file 7m6u at 2.59A resolution of what is now called carboxypeptidase G2 (Yachnin et al., 2022), and it has an Asp-*cis*-Asp in the equivalent position, where the first Asp binds both Zn^++^. With a closely related structure but no detectable overall sequence identity, the 8vkt 1.4A structure of DapE (Terrazas-Lopez et al., 2024) has an Asp-*cis*-Met in the equivalent local position, where the Asp binds both Zn^++^.

### Cys-*cis*-Cys: vicinal disulfides

The *cis*-nonPro list includes two examples of Cys-*cis*-Cys, that are SS-bonded as sequence-adjacent, or “vicinal”, disulfides. Figure 7 shows the *cis* vicinal SS in 1wd4, where the disulfide makes van der Waals contact with bound arabinofuranose. The *trans* conformation is also possible, and more frequent, for vicinal disulfides. Therefore, we have analyzed the occurrence patterns, the possible conformations (2 *cis* and 2 *trans*), and the varied functional roles of vicinal SS in a separate paper (Richardson et al., 2017). They can bind ligands (usually the undecorated side of a ring, as in Fig. 7), stabilize structure, or provide the switch for a large conformational change.

**Figure 7.**
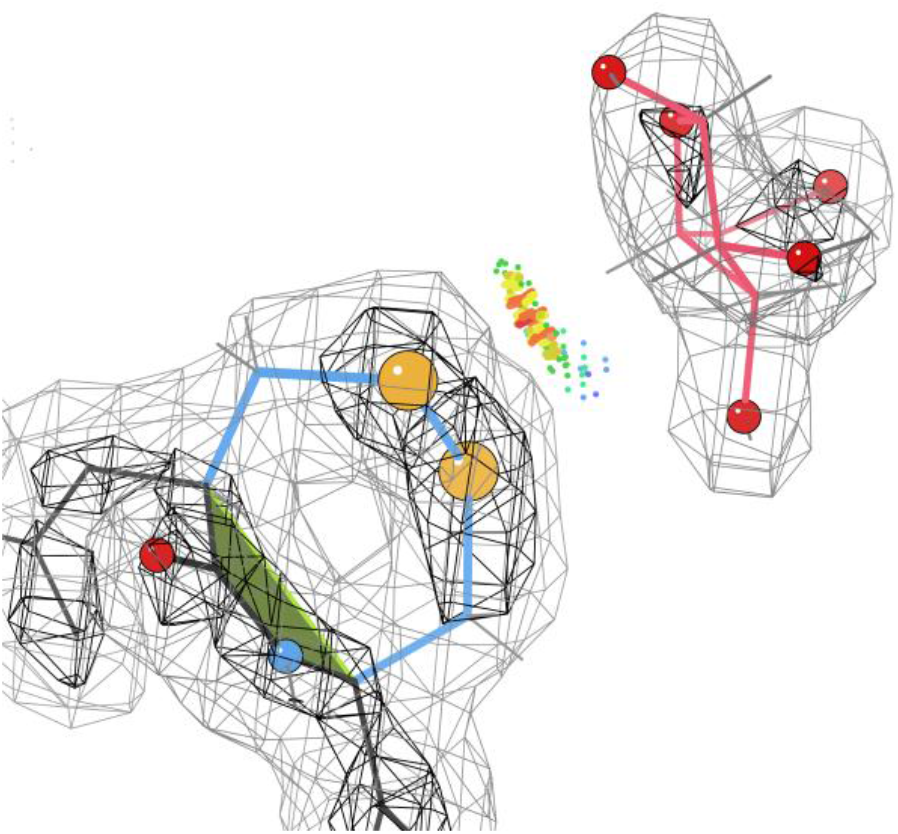
The 1wd4 arabinofuranosidase at 2.04A resolution (Miyanaga et al., 2004). The vicinal SS bond makes extensive van der Waals contact with undecorated side of the sugar ring, preventing binding of a decorated ring. The atoms of the SS bond are gold spheres, and the N and O of the *cis* bond are blue and red spheres, both pointed in the same direction. Green dots are close contacts and orange spikes are small overlaps. Electron density contours are at 1.0 and 3.0σ.

## Discussion

The most novel and striking conclusion of this study is also very puzzling. Why are *cis*-nonPro peptides selected by evolution 4 times more often in carbohydrate-active enzymes than in other proteins, and multiple *cis*-nonPro 36 times as often? As far as we have seen, the *cis* peptide never itself performs the catalysis, and it seems to position a catalytic sidechain or to bind the carbohydrate by many quite distinct strategies. Only a subset of CAZy enzymes, and very few other enzymes, take advantage of these *cis*-nonPro capabilities. Our best guess is that there could be different transition states in carbohydrate chemistry (which is distinct from protein and nucleic acid chemistry) that can make use of the change to a *trans* state to promote more favorable binding (Du et al., 2024). This could begin to be investigated by seeing whether there are any structures of a transition state for these enzymes. There must also be non-proline peptide isomerases that have been coupled into the systems that produce and control carbohydrate-related enzymes. A great many prolyl-peptide isomerases are known, but apparently a generic peptide *cis-trans* isomerase activity has so far been described only for DnaK (Schiene-Fischer et al., 2002), and not with a carbohydrate connection. A search for such enzymes might well be productive and informative.

To support their biological functionality, some *ci*s-nonPros position both sidechains (see Figures 6, 7), while some position just one sidechain (Figures 2b, 5a). Other examples use interactions with the NH and/or CO of the *cis* peptide itself (Figures 1, 3), and those interactions can be either direct or through a water. The two sidechains and the functional groups of the peptide are approximately on opposite sides of the motif. *Cis*-nonPro almost always occur in well-ordered regions of the structure, usually with one side open for business and the other side held by tight contacts, so a *cis*-nonPro in a partly disordered loop is highly suspect. One exception is vicinal disulfides, which can occur and function on very mobile loops, since the disulfide makes a *cis-trans* transition extremely difficult (Richardson et al., 2017). Two situations which have always been found to be wrong are those at chain ends and those that are two-in-a-row (Richardson et al., 2022).

As usually true for rare motifs, nearly all genuine *cis*-nonPro are functionally important. They are rare because they are energetically quite unfavorable relative to *trans*, so they are not conserved by evolution unless they are useful and needed. Of many dozens of examples examined, only a few had no evident connection with function, such as the 4^th^ one in the all-β domain of chitinase 3wd0 and His-*cis*-Ala 759 in 2eab. A major motivation for avoiding or fixing incorrect *cis*-nonPro peptides is to improve the signal-to-noise ratio for recognizing the genuine ones and investigating their functional roles.

## Methods

The starting dataset of *cis*-nonPro examples is from a version of the “Top8000” that includes only the best chain in each RCSB PDB 50% homology cluster, requiring deposited diffraction data, and satisfying other quality criteria. Importantly, the dataset is further quality-filtered at the residue level, aiming to cull out incorrect or unjustified cases and leave only genuine *cis*-nonPro examples. It removes all residues with any backbone atom (including Cβ) with B-factor >40, real-space correlation coefficient <0.7, 2mFo-DFc map value at atom position <1.2σ, a covalent geometry outlier >4σ, an all atom clash ≥0.5 A, or an alternate conformation in the backbone. We could not provide the huge residue-filtered dataset at the time when we did the initial work. However, a later and larger dataset with residue-level filtering is now freely available on Zenodo at https://doi.org/10.5281/zenodo.4626149 (Williams et al., 2022a).

For the *cis*-nonPro work, in addition to automated filtering, 127 cases were examined in 3D along with their electron density, validation markup, and literature references, which identified only two clearly incorrect examples. Both contained a Gly, which in hindsight was reasonable because the lack of a Cβ to provide extra density for a cross-check increases the likelihood of a *trans*-to-*cis* misfit. After finding this, all 84 additional cases were examined that contained a Gly, finding 6 cases to be inadequately supported, which were also removed, and a 4:1 ratio of Gly-*cis*-X to X-*cis*-Gly. All but one were deposited during the period of epidemic over-use of *cis*-nonPro, from 2006-2015. Examination of individual examples was done in the KiNG display and modeling program (Chen 2009), using 2mF_o_-DF_c_ electron density maps and this time always difference-density maps, and their occurrence on partially disordered loops or where identical chains are not *cis* were also rejected.

All figures except Figure 4 were produced in KiNG. The Mage graphics program (Richardson et al., 2001) was used for the 5-dimensional dihedral-angle analysis, because of its cluster-defining functionality and because it can support the 52 lists per subgroup needed to select first- and second-residue sequence and conformation combinations for study.

Occurrence frequencies for the amino acids in *cis*-nonPro peptides were normalized relative to expectation by comparing them with frequencies in the general protein sequence population, as given in the UniProt knowledge base at http://www.ebi.ac.uk/uniprot/TrEMBLstats, version as of November 2017. As described in the text, carbohydrate-active enzymes were identified by their inclusion in the CAZy database at http://www.cazy.org, accessed over mid to late 2016.

In the text and the Supplemental table, a convention is used that is legible in any font (regardless of 0-O, I-1-l confusions): all lower case except for L.

## Supplementary Material

A data table of all genuine *cis*-nonPro cases used here, with PDB code, both residue numbers and amino acids, resolution, number of *cis*-nonPro, abbreviated code, whether in CAZy, tertiary structure, molecule name, notes, CAZy family, and omega:

Table S1.cnP_filterTrue_sort-Cazy-fam.xlsx.

## Acknowledgements

This work was supported by National Institutes of Health grants P01-GM063210 Project IV to JSR and R01-GM073919 to DCR, and the Phenix Industrial Royalties. The authors declare no conflict of interest.

JSR: conceptualization (lead), data curation, formal analysis (lead), funding acquisition, investigation, supervision (lead), validation (lead), visualization (lead), writing – original draft (lead), writing – review and editing. LLV: data curation (lead), formal analysis, investigation (lead), resources (lead), visualization, writing – review and editing. CJW: conceptualization, formal analysis, investigation, software (lead), validation, writing – original draft, writing – review and editing (lead). BJH: formal analysis, investigation, software, writing – review and editing. SML: conceptualization, resources, writing – review and editing. DCR: conceptualization, funding acquisition, project administration (lead), software, supervision, visualization, writing – review and editing.

